# Cannabis Use and Glutamate across the Psychosis Spectrum: In Vivo Evidence from 7T Proton Magnetic Resonance Spectroscopy

**DOI:** 10.1101/2025.10.07.680987

**Authors:** David R. Roalf, Tyler M. Moore, Jacquelyn Stifelman, Maggie K. Pecsok, Ally Atkins, Monica E. Calkins, Mariella De Biasi, Christian Kohler, Christina Mastracchio, Arianna Mordy, Heather Robinson, Ravinder Reddy, Ravi Prakash Reddy Nanga, Kosha Ruparel, Sage Rush-Goebel, Daniel H. Wolf, Ruben C. Gur, Raquel E. Gur, J. Cobb Scott

## Abstract

Cannabis use is linked to elevated psychosis risk, yet the neurobiological mechanisms that couple use to symptom expression remain unclear. Because glutamatergic dysregulation has been implicated in both cannabis effects and psychosis vulnerability, we examined whether brain glutamate relates to dimensional symptoms as a function of cannabis use across the psychosis spectrum. Seventy-nine participants—typically developing controls, clinical high-risk individuals, and patients with psychosis—completed dimensional clinical assessments, detailed cannabis surveys, urine toxicology, and ultra-high-field 7T ^1^HMRS quantification of anterior cingulate cortex (ACC) glutamate levels. Linear models assessed the main and interactive effects of ACC glutamate and cannabis use on positive and negative symptoms. Self-reported cannabis use showed strong concordance with urine toxicology. Cannabis use was associated with higher positive and negative symptoms. Independently, higher ACC glutamate predicted greater positive and negative symptoms. Notably, lower glutamate levels were associated with higher positive symptoms in cannabis users. Exploratory analyses suggested interactions for depressive and manic symptoms, indicating that glutamatergic abnormalities may amplify the overall severity of cannabis-related symptoms. Sensitivity analyses revealed lower ACC glutamate in psychosis patients—especially cannabis users—highlighting diagnostic group differences and reinforcing the link between cannabis exposure and glutamatergic dysfunction. These findings implicate ACC glutamatergic dysfunction as a transdiagnostic correlate of symptom burden, particularly in those with psychosis who are cannabis users. Glutamate-targeted interventions and longitudinal designs will be needed to examine causal pathways linking cannabis exposure to psychosis-relevant outcomes.

---

Recent changes in cannabis laws and the rise in nonmedical cannabis use have intensified concerns about the long-term effects of cannabis on mental health, particularly among youth. While cannabis use has been implicated in a range of psychiatric outcomes^1^, accumulating evidence highlights its association with dimensional psychopathology, including positive and negative psychosis-spectrum symptoms^2^ and mood disturbances^3^. Epidemiological studies show that frequent use—especially of high-potency cannabis—and earlier age of initiation are linked to greater risk for adverse outcomes^4–7^, with a dose–response relationship, particularly in psychosis^8, 9^. Cannabis use during adolescence has been associated not only with the onset of psychotic disorders in early adulthood but also with elevated subclinical symptoms and poorer functional outcomes across multiple domains^10, 11^. In clinical high-risk for psychosis (CHR) samples^12–14^, cannabis use is associated with worsened subthreshold symptoms, and among those with established schizophrenia, it may exacerbate symptom severity and increase relapse risk^15–17^. These effects may be partly mediated by cannabis-induced disruption of the glutamatergic system^18, 19^, a key neurochemical pathway implicated in mood, psychosis, and neurodevelopmental disorders. Both cannabis use^18, 19^ and psychiatric disorders^20–24^ have been independently associated with glutamate dysregulation, yet the mechanistic interplay remains poorly understood. Here we begin to address this gap by integrating detailed self-reported cannabis use, urine drug screening, and ultra–high field 7T proton magnetic resonance spectroscopy (¹HMRS) of the ACC to examine glutamate (Glu) alterations associated with cannabis use across a transdiagnostic sample.

The primary psychoactive component in cannabis, Δ9-tetrahydrocannabinol (THC), modulates neurometabolic and neurotransmitter systems—most notably dopamine and glutamate^19, 25^, both central to psychosis. Sustained cannabis use reduces glutamate signaling *via* Cannabinoid receptor 1 (CB1R) activation, and CB1R agonists induce psychosis-like behaviors^26^. Glutamate hypofunction^20,21^ contributes to dopamine dysregulation in schizophrenia, and recent studies have documented glutamatergic metabolite alterations in individuals with or at risk for schizophrenia^20, 22–24^ and depression^27^. However, systematic *in vivo* investigations of glutamatergic alterations associated with cannabis use in humans using ultra-high field MRI are limited, hindering efforts to clarify how cannabis may heighten psychopathological vulnerability and symptoms.

Glutamatergic alterations are central to the pathophysiology of psychosis. Historically, psychosis has been associated with dopaminergic dysfunction due to its link to positive symptoms^28^, but accumulating evidence emphasize the relevance of glutamate dysregulation in psychosis^29^. A growing literature implicates glutamate dysfunction in psychosis, especially in schizophrenia^30, 31^, including data from animal models^32–36^, postmortem studies^37–40^, genetic analyses^41–45^, peripheral biomarkers^46^, and *in vivo* ^1^HMRS^31^. While early ^1^HMRS meta-analyses at 3T reported higher Glu metabolites in cases vs. controls^47^, subsequent mega-analyses^20^, 7T GluCEST investigations^22, 24^, and 7T ^1^HMRS^23^ studies have demonstrated regionally lower Glu in individuals with psychosis, but see^48^. Region-specific effects include lower Glu in mPFC, frontal white matter, medial temporal lobe, and thalamus, with elevated Glu in cerebellum and basal ganglia.^47^ Taken together, ^1^HMRS studies reveal regionally heterogeneous findings in glutamatergic levels in CHR patients to those with threshold psychosis disorder,^30, 49–54^ suggesting that glutamate alterations may be part of illness progression providing significant rationale for studying glutamate across the psychosis spectrum.

Few studies have directly examined cannabis–glutamate associations across clinical dimensions of the psychosis spectrum, and existing work has been limited to 3T MRI. Early ^1^HMRS work found that patients with early psychosis who used cannabis exhibited lower prefrontal Glu concentrations than both non-using psychosis patients and healthy controls, along with a steeper age-related Glu decline, suggesting that cannabis may exacerbate glutamatergic dysfunction and influence disease progression^55^. Another 3T ^1^HMRS study in early psychosis patients observed limited differences in Glutamate+Glutamine (Glx) between cannabis users and nonusers but identified associations between regional gray matter volume (caudate, hippocampus, cortex) and caudate Glx in cannabis users, implying that heavy cannabis use may alter structure–neurochemical connectivity in CB1R-rich regions^56^. In healthy individuals, acute THC induced greater Glx surges in those who developed transient psychotic-like symptoms, particularly among participants with lower baseline Glx and greater prior cannabis exposure^57^. These findings collectively suggest that baseline glutamatergic state and cannabis responsiveness may interact to influence psychosis risk and expression. However, prior studies have been limited to 3T ^1^HMRS, where sensitivity to glutamate is lower, and they have only measured cannabis–glutamate associations in categorical clinical groupings.

In summary, converging epidemiological, preclinical, and neuroimaging evidence implicates glutamate as a key modulator of the cannabis–psychosis relationship. By integrating detailed cannabis use phenotyping, high–field 7T ^1^HMRS measures, and dimensional clinical assessments across individuals who are typically developing, CHR or experience psychosis (PSY), our study aims to elucidate how cannabis use is associated with glutamatergic levels in those with and at risk for psychosis. In line with recent RDoC and NIMH initiatives, we assess clinical symptoms dimensionally rather than relying solely on categorical diagnoses. Dimensional measurement provides greater sensitivity to subthreshold variation^58, 59^, allowing us to capture the full spectrum of psychopathology and potential associations with cannabis use, including effects that may not meet diagnostic thresholds. This approach also facilitates transdiagnostic comparisons and improves power to detect brain–behavior associations by reducing heterogeneity within broad diagnostic categories. Understanding this interplay may identify neurobiological and clinical markers of risk and ultimately inform targeted prevention and intervention strategies for this high-risk population.

## Methods

### Participants

To capture a broad range of psychopathology, we studied both healthy individuals and those with subthreshold and threshold psychiatric symptoms. The study sample (N = 79, ages 14–32, 35 female) included a typically-developing (TD) group (no Axis I diagnoses or subthreshold psychosis symptomatology, no history of psychotropic medication use; N = 31) and a clinical group composed of individuals classified by experienced clinician scientists (CK, MEC) as having psychosis spectrum disorders (N = 48), including 27 with CHR and 21 with schizophrenia spectrum and other psychotic disorders (PSY; N = 21). Although our primary analyses treat psychopathology as a continuous, transdiagnostic construct, demographic and clinical characteristics are displayed by categorical diagnosis for clarity and interpretability. This allows for evaluation of the representativeness of the sample and provides context for the dimensional analyses presented below. Complete demographic and clinical information are presented in Table 1, stratified by cannabis user status and clinical diagnosis. In addition, sensitivity analyses are conducted by diagnostic group (Supplemental Results). Brief cognitive (Montreal Cognitive Assessment (MoCA^60^)) and functional assessments (Global Assessment of functioning (GAF^61^)) were also administered.

**Table 1.** Participant demographics and clinical characteristics. This table summarizes demographic and clinical characteristics of the study sample. Demographic variables include age, sex distribution, and relevant clinical status or group designation. Mean (+/- s.d.) positive, negative, disorganized and generalized SIPS, GAF and MoCA scores are shown.

| DxGroup |  | N | Age<br>(years) | Percent<br>Female | SIPS<br>Positive | SIPS<br>Negative | SIPS<br>Disorganized | SIPS<br>Generalized | GAF | MoCA |
| --- | --- | --- | --- | --- | --- | --- | --- | --- | --- | --- |
| **TD** |  |  |  |  |  |  |  |  |  |  |
| **Cannabis User** |  |  |  |  |  |  |  |  |  |  |
|  | No | 16 | 20.7 ± 2.7 | 63% | 1.3 ± 1.8 | 1.1 ± 1.0 | 0.4 ± 0.6 | 0.7 ± 0.9 | 87.4 ± 4.5 | 26.5 ± 2.8 |
|  | Yes | 15 | 22.8 ± 3.0 | 27% | 1.7 ± 1.8 | 1.5 ± 1.8 | 0.3 ± 0.6 | 1.5 ± 2.0 | 81.2 ± 13.7 | 27.7 ± 1.6 |
| **CHR** |  |  |  |  |  |  |  |  |  |  |
| **Cannabis User** |  |  |  |  |  |  |  |  |  |  |
|  | No | 14 | 20.8 ± 3.6 | 50% | 8.1 ± 4.7 | 7.6 ± 6.0 | 4.1 ± 2.5 | 3.7 ± 3.4 | 56.9 ± 8.0 | 24.8 ± 2.9 |
|  | Yes | 13 | 22.6 ± 2.8 | 31% | 6.8 ± 3.1 | 7.6 ± 5.5 | 4.0 ± 3.0 | 3.2 ± 2.5 | 61.0 ± 12.2 | 27.0 ± 2.7 |
| **PSY** |  |  |  |  |  |  |  |  |  |  |
| **Cannabis User** |  |  |  |  |  |  |  |  |  |  |
|  | No | 5 | 26.8 ± 4.1 | 60% | 14.0 ± 6.1 | 9.4 ± 5.6 | 5.4 ± 1.1 | 5.6 ± 3.6 | 47.4 ± 4.0 | 22.0 ± 3.9 |
|  | Yes | 16 | 24.8 ± 3.5 | 44% | 19.7 ± 6.0 | 12.0 ± 2.7 | 7.6 ± 4.2 | 8.5 ± 6.0 | 42.4 ± 9.0 | 25.6 ± 2.4 |

### Clinical Assessment

All participants received detailed clinical assessments, including a semi-structured diagnostic interview to assess a broad spectrum of psychosis-relevant experiences, psychopathology, treatment history, and medications. Participants diagnostic assessments by highly-trained assessors using a computerized version of a modified and adapted *Schedule for Affective Disorders and SZ for School Age Children – Present and Lifetime Version (K-SADS-4.0)*^62^, which includes semi-structured assessments of mood, ADHD and substance use disorders. Psychosis spectrum symptoms were assessed using the *Structured Interview for Prodromal Syndromes (SIPS, v. 4.0)*, in which selected modules of the Structured Clinical Interview for DSM-IV (SCID-IV)^63^ are integrated to facilitate differential diagnosis. The *Scale of Prodromal Symptoms (SOPS)*^64^, embedded within the SIPS^65^, describes and dimensionally rates the severity of positive, negative, disorganized, and general symptoms occurring within the past 6 months. To facilitate efficient data collection, the instruments were integrated into one administrative tool, the Computer-Assisted Psychopathology Assessment (CAPA)^62^. Interviewers underwent formal training including lectures and mock interviews, followed by supervised administration and performance evaluation until competency was established^62^. For minors, collateral information was provided by an independent interview with a parent or guardian. All information was integrated to yield consensus diagnoses and clinical symptom ratings at a clinical case conference led by at least two doctoral-level clinicians. CHR was defined as the presence of at least one positive symptom rated 3-5, or at least two negative and/or disorganized symptoms rated 3-6. Psychosis (PSY) patients are individuals with a DSM-5 schizophrenia spectrum disorder diagnosis (e.g., SZ). Clinical scores are shown in Table 1.

### Estimation of Correlated Traits Clinical Factor scores

Clinical factor scores were derived using a framework that incorporated longitudinal computerized assessments (e.g. GOASSESS^66, 67^ and CAPA^68^) used in our laboratory group since the landmark Philadelphia Neurodevelopmental Cohort study.^69–71^ We used a bank of all clinically available data from our laboratory to inform a dimensional factor analysis (e.g. GOASSESS^66, 67^ and CAPA^68^) and generate clinical factor scores for participants enrolled in the current study.

Because the same instruments were not used across historical time points, historical data were harmonized across longitudinal visits by using only items that overlapped between GOASSESS and CAPA interviews. These included the screen items from the depression, mania, and ADHD sections of the GOASSESS/CAPA, as well as the PRIME^72, 73^ screen and a subset of SIPS items. Using this harmonized framework, dimensional clinical scores (correlated trait factor scores) were generated for each participant in the present study based on the CAPA data collected (described in the Clinical Assessment section). Dimensional clinical factors scores are shown, by diagnostic group, in Supplemental Table 2.

### Cannabis Use

Cannabis use was measured using a computerized version of the Minnesota Center for Twin and Family Research self-report substance use assessment^74^, which was privately self-administered on a computer. The measure assessed lifetime use of several substances; for cannabis and alcohol, additional questions queried age at first use and frequency of past year use. To obtain more detailed data regarding patterns of cannabis use, several questions from the *DFAQ-CU* (*Daily Sessions, Frequency, Age of Onset, and Quantity of Cannabis Use Inventory*)^75^ were also administered. This measure provided data for individual methods of cannabis use, with visual depictions to indicate quantities of cannabis across forms (e.g., joint, flower). Summary measures were generated for frequency, age of onset, and quantity of cannabis use across methods of use, including increasingly popular methods (e.g., edibles, concentrates). The DFAQ-CU shows convergent, predictive, and discriminant validity, including high expected correlations (>0.7) with other frequency measures (e.g., timeline follow-back), modulates correlations with cannabis use disorder symptoms and cannabis-related problems, and low correlations with alcohol use problems^75, 76^. Consistent with prior research^77, 78^, we classified individuals as *<u>cannabis users</u>* if they reported use more than once per month over the past year; for exploratory analyses we also classified individuals as *<u>frequent cannabis users</u>* if they reported using cannabis on a daily or nearly daily basis for the past 12 months. We also administered sections of the *Cannabis Experiences Questionnaire (CEQ)* that assess motivations for cannabis use and positive and negative experiences associated with use, consistent with prior studies in early psychosis^79^.

### Urine Drug Screen for Cannabis

In line with expert consensus groups recommendations^80^, we also used a urine drug test to identify presence/absence of cannabis use. All urine samples (90ml) were collected using Redwood Toxicology iCUPs™, which detect one or more of the primary metabolites of THC. The immunoassay is binary (positive or negative). If the sample is below threshold, it is reported as negative; if above, it is reported as positive. Detection threshold for presence/absence is 50 ng/mL of THC-COOH. Cannabis metabolites are typically detected for up to 3 days after a single use in occasional users, while frequent users typically test positive for a much longer period—sometimes up to several weeks—due to the slow release of stored THC from fatty tissues and repeated, high-level exposure. Body fat percentage, metabolism, hydration level, and dosage all affect how long THC-COOH remains detectable in urine. Urine drug screening occurred prior to the 7T MRI session. If a participant reported cannabis use on the day of the MRI, or if acute intoxication was suspected, the study visit was rescheduled.

### 7T ^1^HMRS Data Acquisition

All ^1^HMRS data were acquired on a whole-body 7 Tesla Terra MRI system (Siemens, Erlangen, Germany) equipped with a single-channel volume transmit and 32-channel receive proton phased array head radiofrequency coil (Nova Medical, Wilmington, MA, USA). Prior to spectroscopy, high-resolution structural images were obtained using a magnetization-prepared rapid acquisition gradient echo (MP2RAGE) sequence (TE = 2.52 ms, TR = 5000 ms, TI1 = 700 ms, TI2 = 2500 ms, flip angle 1 = 7°, flip angle 2 = 5°, voxel size = 0.8 mm^3^). These anatomical images served both for voxel placement and for later tissue-segmentation corrections.

### Voxel Localization and Shimming

^1^HMRS voxels (10 x 40 x 15 mm³) were manually placed in anterior cingulate cortex, using the mid-sagittal anatomical image as reference. Care was taken to align all voxel faces orthogonal to the cortical surface and to minimize inclusion of cerebrospinal fluid (CSF)-rich sulcal regions. First- and second-order B₀ shimming was performed manually over each voxel to achieve a full-width at half-maximum (FWHM) linewidth of ≤ 20 Hz for the unsuppressed water peak within the voxel.

### Spectroscopy Sequence

Spectra were acquired using a point-resolved spectroscopy (PRESS) sequence optimized for 7T. Sequence parameters were as follows: TE = 23 ms, TR = 3,000 ms, number of averages (NA) = 64 (total acquisition time ≈ 3 min 24 s per voxel). Unsuppressed water reference scan preceded this acquisition for reference and calibration to quantify and correct for eddy-currents. Sequence parameters of the unsuppressed scan (TE = 23 ms, TR = 3,000 ms, NA = 8) used identical localization and shimming settings.

### Data Processing and Quantification

All spectroscopy data were exported in RDA format, which were coil-combined and averaged at the scanner. All spectra were processed, modeled, and quantified using Osprey (v2.9.0; Osprey Project, Johns Hopkins University)^81^ running in MATLAB (R2023b; MathWorks, Natick, MA, USA). Spectral fitting was performed using the LCModel package (v6.3-1N) implemented in Osprey software. Prior to fitting, imported raw spectra were processed through the ‘OspreyProcess’ module, which includes eddy-current correction, frequency and phase alignment, water removal, frequency referencing, and initial phasing. Default parameters were utilized for modeling and quantification in ‘OspreyFit’, including a metabolite fit range of 0.5 to 4.0 ppm, a water fit range of 2.0 to 7.4 ppm, and a knot spacing of 0.4 ppm. The basis set provided for LCModel included simulated metabolite spectra for N-acetylaspartate (NAA), creatine (Cr), choline (Cho), glutamate (Glu), glutamine (Gln), myo-inositol (mI), γ-aminobutyric acid (GABA), and others, generated with density-matrix simulations at 7T incorporating sequence-specific TE and TM. In addition to the basis set, default macromolecular and lipid components provided by Osprey were fitted to each spectrum. Each participant’s ^1^HMRS voxel was then co-registered to a T_1_-weighted MP2RAGE provided for each individual. Co-registration in ‘OspreyCoReg’ also produces a voxel mask that is subsequently used in ‘OspreySeg’ to segment the voxel into gray matter, white matter, and CSF using SPM12. Example spectra and voxel overlap across participants are shown in Figure 1. In this module, fractional tissue volumes are also determined. Tissue- and relaxation-corrected metabolite molal concentrations were then estimated in ‘OspreyQuantify’ according to the Gasparovic method^82^. Only tissue-corrected metabolite estimates passing quality control (Cramér–Rao lower-bound (CRLB) ≤ 20%) were included in subsequent analyses.

**Figure 1.**
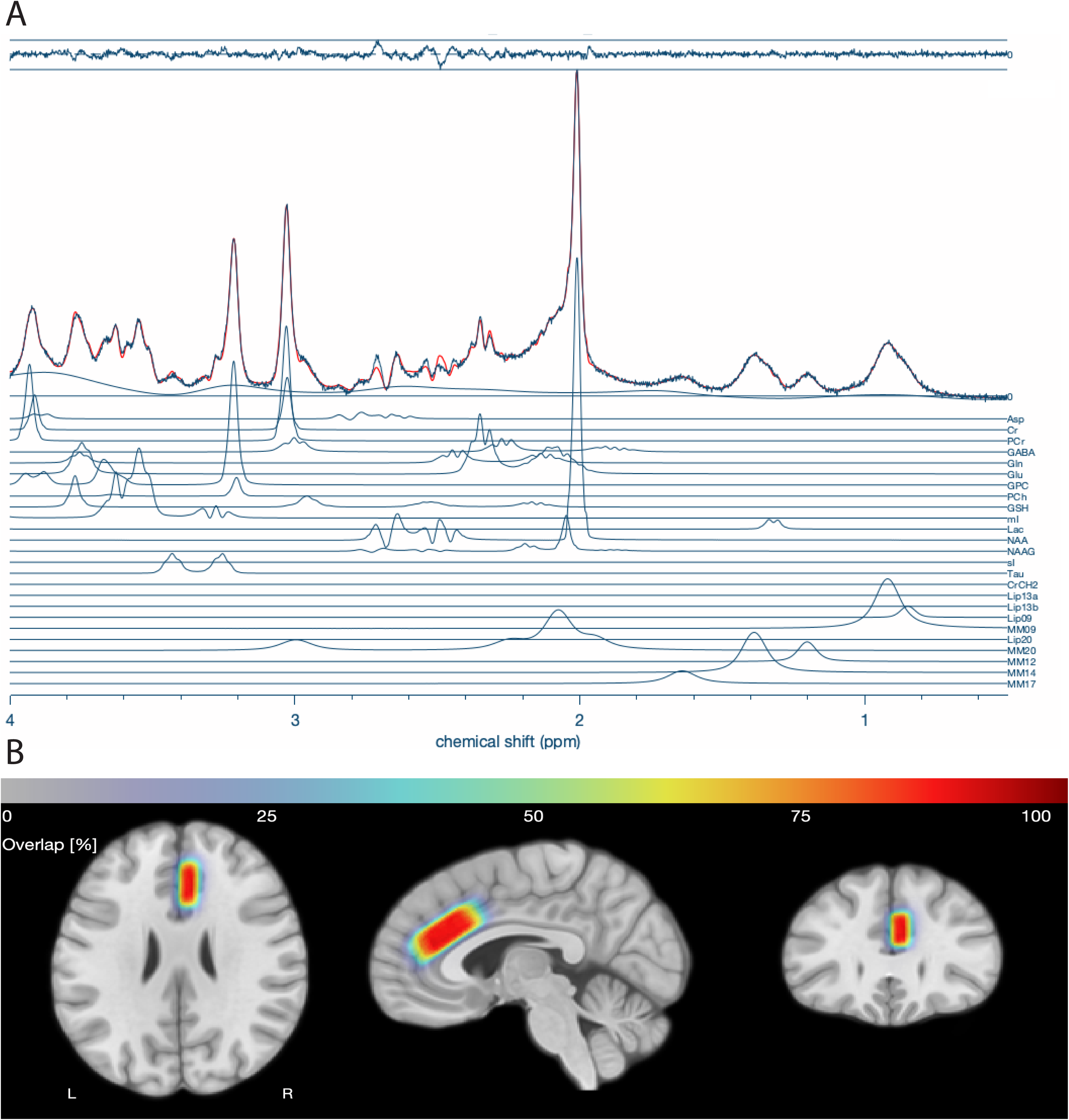
Individual metabolite quantification and voxel overlap in single-voxel spectroscopy (SVS) ¹H-MRS. Panel A displays individual metabolite estimates obtained using the Osprey analysis pipeline. Each line represents the concentration of a specific metabolite quantified within the spectroscopy voxel for a representative participant. Key metabolites include glutamate (Glu), γ-aminobutyric acid (GABA), N-acetylaspartate (NAA), creatine (Cr), choline (Cho), myo-inositol (mI), and others derived from short-TE PRESS acquisitions. Panel B shows a voxel overlap heatmap across all participants, generated by co-registering individual SVS voxel placements to a common template space. Warmer colors indicate regions where greater numbers of participants’ voxels overlap, demonstrating the consistency of voxel localization across the sample and providing a measure of spatial coverage and reproducibility of the MRS acquisition.

### Statistical Analysis

Group differences in cannabis use rates were assessed using independent-samples t tests and chi-squared tests, as appropriate. To examine the effects of ^1^HMRS glutamate, cannabis use, and their interaction on clinical symptom dimensions, we conducted a series of linear regression models. The primary outcome measures of interest were those associated with psychosis: positive and negative symptoms. Each of the measures was modeled as a continuous outcome. All linear models controlled for sex and age at scan. Cannabis use was coded dichotomously (0 = non-user, 1 = user), and glutamate was entered as a continuous variable. Interaction terms between glutamate and cannabis user status were included to assess modulation effects. Significance was assessed at *p* < 0.05, with trend-level effects noted at *p* < 0.10. Residuals were visually inspected for normality and homoscedasticity. Results are reported as F-statistics with associated degrees of freedom and *p*-values. Exploratory analyses in other symptom domains (depression, mania, and ADHD) were also conducted. As a sensitivity analysis, these analyses were repeated with categorical diagnostic group (TD, CHR, PSY; See Supplemental Results). Demographic data were compared across diagnostic groups (e.g., diagnosis, cannabis user status) using t-tests and chi-square as appropriate. Rates of cannabis use and urine drug screen data were compared across all three diagnostic groups using an omnibus Chi-square test, with planned post-hoc pairwise comparisons. Post-hoc comparisons were completed using simple slopes (‘sim_slopes’) and least squares means approach (‘lsmeans’) using the ‘interactions’^83^ and ‘lsmeans’^84^ libraries in R (V4.3.3), respectively.

## Results

### Self-reported cannabis use rates converge with results of urine toxicology screening

Across the sample, 56% (n=44) self-reported lifetime cannabis use, including 32% (n=25) frequent users and 24% (n=19) infrequent users, while 44% (n=35) were cannabis non-users (Figure 2). To assess the validity of self-reported cannabis use, we compared responses on cannabis use self-report assessments to results from urine toxicology across all individuals. 89% (70/79) of individuals had a valid urine toxicology screen. A chi-square test revealed a significant association between self-reported use and toxicology results, χ²(1) = 7.62, *p* < 0.01. The vast majority (92%) of cannabis non-users tested negative on the urine toxicology screen (Figure 3A), while self-reported cannabis users were more likely to test positive on the urine toxicology screen (Figure 3B), but overall rates of positive screens were low (38%), which is not unexpected given the typical window of cannabis metabolite detection. Indeed, self-reported frequent users (87%) were more likely to test positive on urine toxicology than infrequent users (13%; Figure 3C). Overall, these results indicate good concordance between self-report and biological verification of cannabis use. Note that two individuals who denied cannabis use tested positive on urine toxicology; these individuals were considered infrequent cannabis users in subsequent analyses.

**Figure 2.**
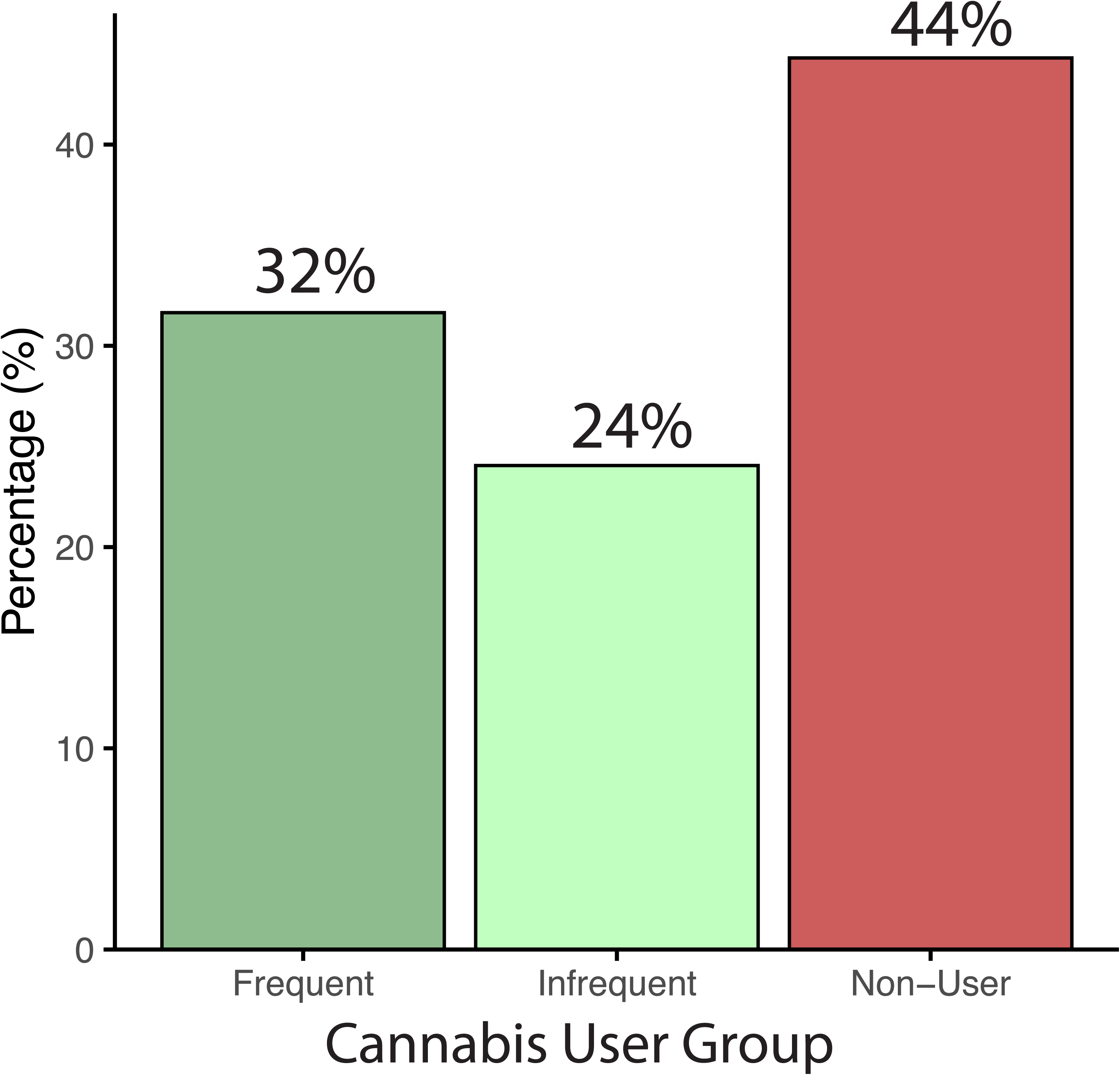
Frequency of cannabis use in the transdiagnostic sample who underwent 7T MRI.

**Figure 3.**
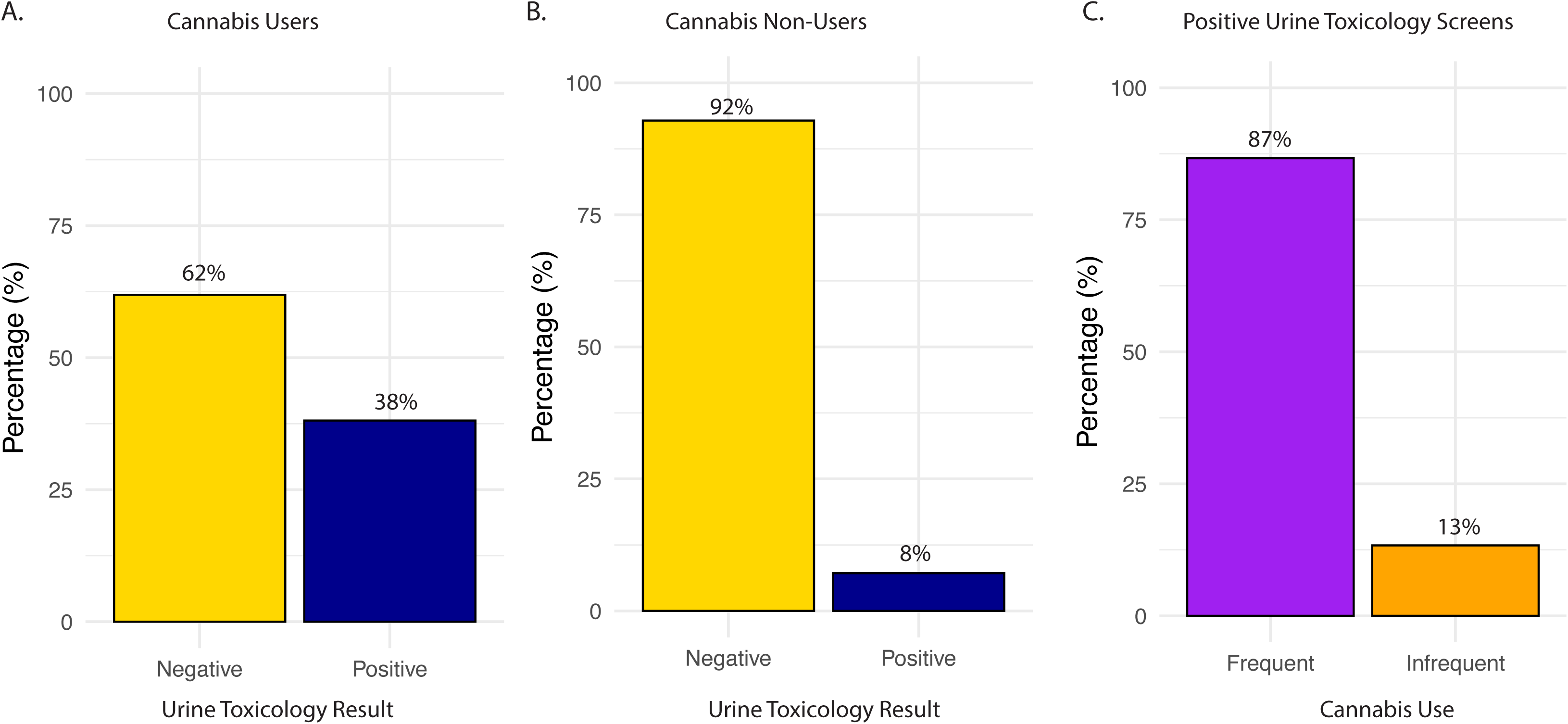
Comparison of self-reported use and urine toxicology screening. A. Many, but not a majority, of self-reported cannabis users tested positive on a urine toxicology screen. *Note two cannabis users did not complete urine toxicity screening. B. The vast majority of self-reported cannabis non-users tested negative on a urine toxicology screen. *Note seven cannabis non-users did not complete urine toxicity screening. C. Self-reported frequent cannabis users were more likely to test positive on a urine toxicology screen as compared to infrequent users.

### Cannabis use is higher in psychosis patients

76% of PSY participants, 48% of CHR participants, and 45% of TD participants reported cannabis use (Supplemental Figure 1). A chi-square test of independence comparing all three groups was not significant (χ²(2) = 5.51, *p* = 0.06), but planned pairwise comparisons indicated that cannabis use was more prevalent in the PSY group compared to the TD group, χ²(1) = 3.78, *p* = 0.05. The rate of use between PSY and CHR groups showed nominally more cannabis use in PSY (χ²(1) = 2.80, *p* = 0.09), but CHR and TD groups did not differ in cannabis use, χ²(1) < 0.01, *p* = 0.99. Notably, analysis of frequency of cannabis use (Supplemental Figure 2) in CHR (33%) and PSY (32%) showed similar rates of frequent use (3-4 times a week or daily).

### Glutamate and cannabis use are associated with dimensional clinical symptoms

We examined the relationship of ^1^HMRS glutamate levels, cannabis use, and their interaction with dimensional clinical symptoms across the entire sample. A significant Glu × cannabis interaction was observed for positive psychosis symptoms (*F*(1,68) = 4.33, *p* = 0.041), indicating that the relationship between glutamate and positive symptom severity differed by cannabis use status (Figure 4A). Using a simple slopes follow-up analysis, in cannabis users, lower glutamate was significantly associated with greater positive symptom severity (β = –0.15, *SE* = 0.05, *t*(68) = –3.11, *p* = 0.003), but this association was not significant in non-users (β = 0.01, *SE* = 0.06, *t*(68) = 0.23, *p* = 0.82). These findings suggest that glutamate levels are differentially linked to positive psychosis symptoms depending on cannabis use history, with effects most pronounced among cannabis users.

**Figure 4.**
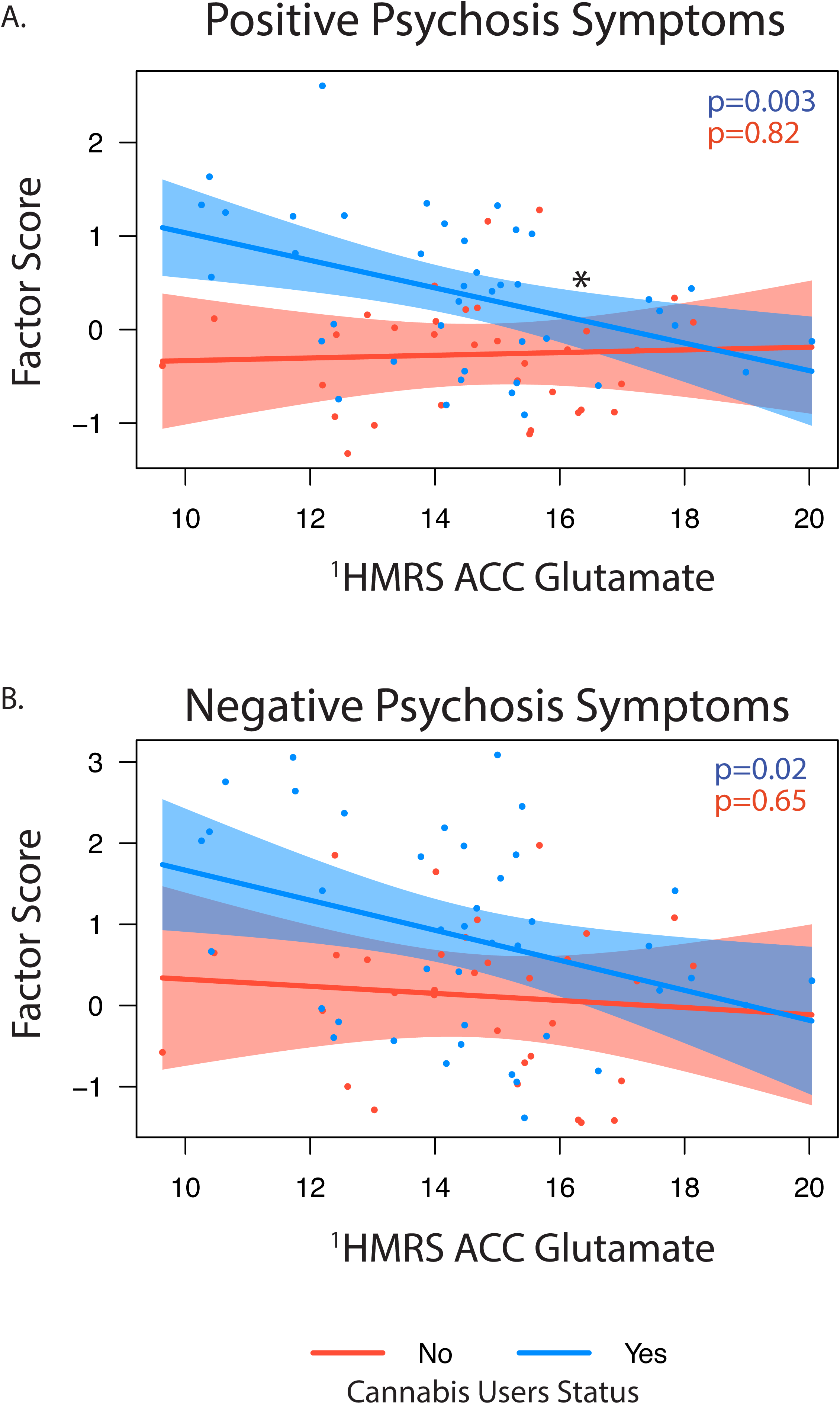
Associations between ^1^HMRS glutamate level and positive (A) and negative (B) psychosis factor scores stratified by cannabis use status (yes/no). Simple slope significance (p) values are show for each association. *indicates significant interaction.

Additionally, both glutamate (*p* = 0.012) and cannabis use (*p* < 0.001) independently predicted greater positive symptoms. For negative psychosis symptoms (Figure 4B), glutamate (*p* = 0.01) and cannabis use (*p* = 0.02) were also significant predictors, but there was no interaction, suggesting additive effects of elevated glutamate and cannabis exposure. Follow-up sensitivity analysis comparing glutamate levels by psychosis diagnostic group showed similar results, including when stratified by cannabis use status (Supplemental Results).

In the exploratory analysis of other dimensional symptoms, cannabis use was associated with greater depressive symptoms severity (*p* = 0.011), and a trend-level interaction with glutamate (*p* = 0.05), which suggested a potential cannabis-dependent relationship between glutamate and depressive symptoms (Figure 5A). There was no main effect of glutamate (*p* = 0.22). A similar pattern emerged for manic symptoms, where cannabis use again predicted greater symptom burden (*p* = 0.01), while glutamate showed a marginal association (*p* = 0.09), but with no significant interaction (Figure 5B). In contrast, ADHD symptoms were not significantly associated with either glutamate or cannabis use (Figure 5C). Higher glutamate levels were also associated with overall functioning as measured by the GAF (F(1,66)=4.50. *p* = 0.03), with cannabis non-users showing trend-level (*p* = 0.06) higher GAF scores than cannabis users. Within the cannabis user group, there were no associations between glutamate level and age of onset, length of use, or paranoia scores on the CEQ. There was a trend level association for cannabis-associated euphoria, with lower levels of glutamate associated with higher reported CEQ euphoric scores (*p* = 0.07). Analyses stratified by clinical diagnosis showed similar trends (Supplemental Results).

**Figure 5.**
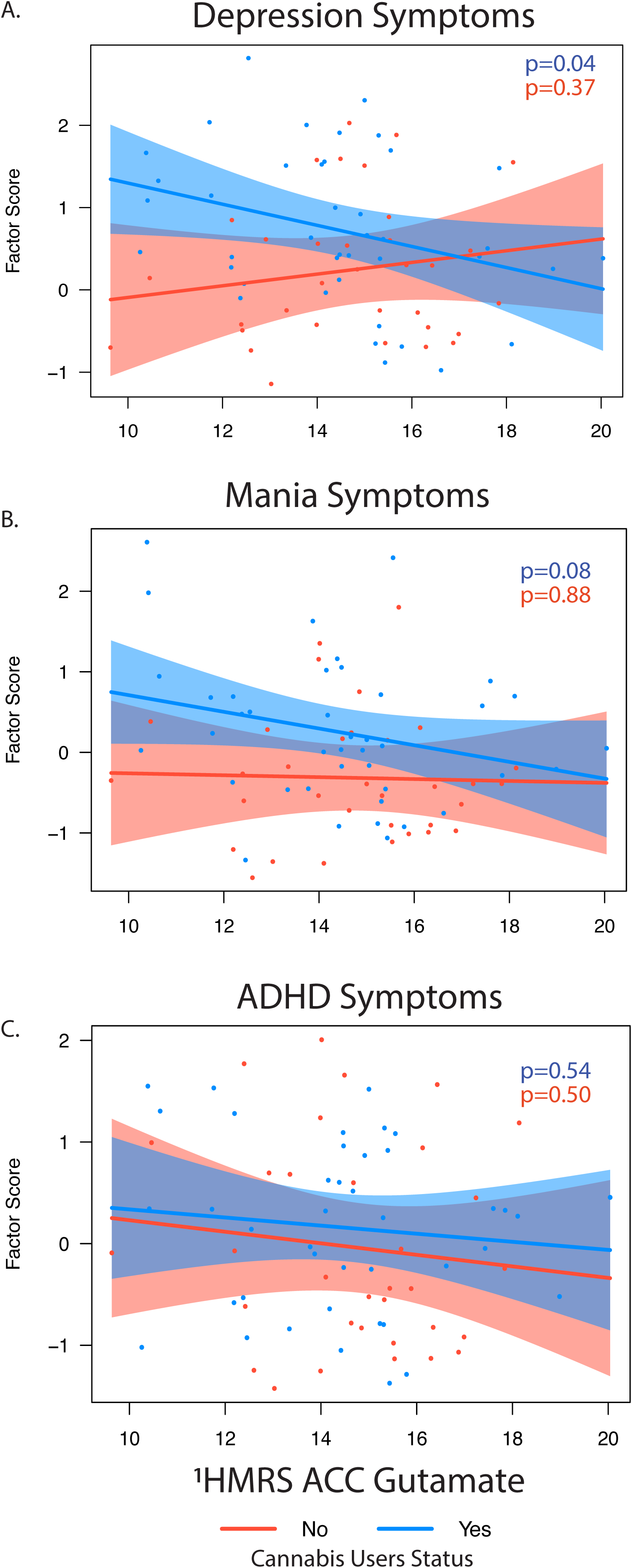
Exploratory associations between ^1^HMRS glutamate level and depression (A), mania (B) and ADHD (C) factor scores stratified by cannabis use status (yes/no). Simple slope significance (p) values are show for each association. *indicates significant interaction.

## Discussion

We combined high-resolution 7T ^1^HMRS with detailed cannabis use assessments to examine glutamatergic alterations in a transdiagnostic sample of patients with psychosis, clinical high risk for psychosis, and typically developing youth. Cannabis use was common overall but most prevalent among individuals with psychosis. Self-reported use showed strong concordance with urine toxicology results, especially among frequent users, supporting the validity of self-report in this cohort. As expected, psychosis participants exhibited elevated clinical symptomatology and lower global functioning compared to CHR and TD groups. Collectively, the results indicate that cannabis use is robustly associated with elevated symptoms across multiple clinical domains—particularly psychosis, but also depression and mania—and that glutamate levels contribute independently to symptom severity, with interaction effects most apparent for positive symptoms, and potentially depressive symptoms. These findings suggest that glutamatergic dysregulation may play a role in symptom expression among cannabis users, especially for the positive symptom dimension of psychosis.

A novel finding of the current study was that anterior cingulate glutamate levels were significantly associated with positive psychosis symptoms only among cannabis users, suggesting that cannabis use modulates the glutamate–psychosis relationship. Specifically, lower glutamate predicted greater positive symptom severity in cannabis users but not in non-users. This pattern is consistent with evidence that cannabis can disrupt glutamatergic signaling in CB1 receptor–rich cortical regions, potentially leading to maladaptive reductions in excitatory tone or altered glutamate homeostasis in circuits already vulnerable to dysregulation in psychosis^85^. Such cannabis-related alterations could diminish glutamatergic efficiency, thereby linking lower glutamate levels to greater positive symptom expression among users. These results raise the possibility that ACC glutamate could serve as a biomarker of vulnerability among cannabis-using individuals on the psychosis spectrum, which could inform risk stratification and targeted interventions.

Importantly, glutamate was also independently associated with negative symptoms and global functioning across groups, and cannabis use predicted worse outcomes on these measures, indicating broader relevance of glutamatergic dysfunction beyond cannabis-exposed individuals. Given prior reports of regionally specific glutamate reductions in early and chronic psychosis^22, 23, 85^, the additive and interactive effects of cannabis use observed here may reflect either direct neurochemical consequences of cannabis or shared neurodevelopmental risk factors, or a combination of both. These findings support a mechanistic framework in which cannabis use interacts with regional glutamatergic vulnerability to influence the severity and profile of psychopathology. Future longitudinal and ultra–high field imaging studies will be critical to further clarify causal pathways linking cannabis use, glutamate dysregulation, and clinical outcomes, and to determine whether glutamate-targeted pharmacological or behavioral interventions may be particularly beneficial for cannabis-using individuals with psychosis.

We also found that cannabis use was associated with greater severity of depression and mania symptoms across the whole sample, highlighting the transdiagnostic clinical relevance of cannabis use. This is consistent with 1) meta-analytic evidence that heavy cannabis use increases risk for depressive disorders^86^; and 2) epidemiological findings of high comorbidity between cannabis use and mood disorders^87, 88^. The present associations may reflect overlapping neurobiological vulnerabilities—such as altered serotonergic and endocannabinoid signaling—that predispose individuals to both cannabis use and mood dysregulation, or the downstream effects of cannabis-related perturbations in neurotransmitter systems on affective processing.

Self-reported cannabis use showed high concordance with urine toxicology results, supporting the validity of self-report in assessing cannabis exposure in psychiatric populations. Across the sample, 92% of individuals denying cannabis use tested negative on urine toxicology, and frequent users were far more likely to test positive than infrequent users (87% vs. 13%). This pattern replicates prior work showing that urine toxicology has high specificity and negative predictive value for cannabis^82, 89^ and that concordance is often strongest among individuals with frequent use^90^.

Nonetheless, some discrepancies were observed, with two participants denying cannabis use testing positive on toxicology and overall low rates of positive screens among self-reported users (38%), underscoring important limitations of toxicology measures. Detection windows for urine THC metabolites are finite and may under-detect use in infrequent users^91^, while contextual factors, stigma, and recall error could bias self-reports^92, 93^. These findings highlight the utility of pairing structured self-report cannabis use measures with biological verification, particularly in research on high-risk groups such as individuals with psychosis, where cannabis use is prevalent and clinically relevant^94^. Combining approaches may increase confidence in exposure classification while also capturing use patterns and frequency not discernible from toxicology alone.

Notwithstanding the strengths of combining ultra–high field 7T ^1^HMRS, biological and self-report measures of cannabis use, and dimensional clinical assessment across a transdiagnostic sample, several limitations should be noted. First, the cross-sectional design limits causal inference about the relationship between glutamate levels, cannabis use, and psychopathology symptom expression. Longitudinal studies are needed to determine whether cannabis-related glutamatergic alterations precede or result from symptom exacerbation. Second, although self-reported cannabis use showed strong concordance with urine toxicology results, the binary nature and limited detection window of the urine screen may underestimate recent use, particularly in infrequent users. Third, although the inclusion of clinical high-risk, psychosis-spectrum, and typically developing individuals enhances generalizability, the modest sample size, particularly within diagnostic subgroups, may limit statistical power to detect more nuanced interaction effects or subgroup-specific associations. Fourth, while advanced tissue correction and spectral quality control were employed, ^1^HMRS glutamate measures remain unable to fully disentangle glutamate from glutamine, do not include information on inhibitory neurometabolites (e.g., GABA), and do not reflect synaptic neurotransmission directly. Lastly, potential confounding effects of other substances (e.g., alcohol, nicotine), antipsychotic medication^48^ or comorbid psychiatric conditions were not fully modeled, and future work should expand the analytic framework to include polysubstance exposure and broader transdiagnostic factors.

In conclusion, these findings suggest that cannabis use may interact with psychosis-related vulnerability to accentuate glutamatergic dysfunction, and that this neurochemical alteration may contribute to symptom expression, especially in the domain of positive symptoms. Understanding this relationship is critical to elucidate the neurobiological substrates underlying both acute and chronic cannabis effects on mental health.

## Acknowledgements

We thank the patients and families who participated in this study.

These data, in part, were presented at the 2025 Congress of the Schizophrenia International Research Society in Chicago, IL, USA.

## Availability of data and materials

The data generated and/or analyzed during the current study is available. Please contact <u>Dr. David Roalf</u> with questions and considerations for data sharing upon reasonable requests.

## Author Information

### Contributors Statement

All authors contributed to the writing, editing, and approval of this manuscript. DRR and JCS conceptualized and designed the research. MEC, CK, KR, SGR, RCG, JS, HR, AM, CM and REG, prepared and/or collected data on the assessment tools. DRR, AA, & KR performed aspects of the MRI experiments and analysis. DRR, TMM, KR, & JCS performed the statistical analyses. DRR wrote the first draft of the manuscript. All authors critically reviewed the manuscript’s content and approved the final version for publication.

### Corresponding author

Correspondence to <u>Dr. David Roalf</u>

### Ethics declarations

Ethics approval and consent to participate.

Participants provided informed consent/assent and the study procedures were approved by the Institutional Review Boards at the Children’s Hospital of Philadelphia and the University of Pennsylvania. Participants’ privacy and confidentiality were ensured at every stage of the study. We confirm that all methods were performed in accordance with the relevant guidelines and regulations.

### Consent for publication

Not applicable

