## Supplemental Material for "Cannabis Use and Glutamate across the Psychosis Spectrum: In Vivo Evidence from 7T Proton Magnetic Resonance Spectroscopy"

**Supplemental Methods
Method for Estimation of Correlated Traits Clinical Factor scores.**

Estimation of clinical factor scores was done using an exploratory structural equation model (ESEM) with geomin rotation and clustering by subject due to the repeated-measures nature of the data. The number of factors to extract was determined by interpretability and visual examination of the scree plot (based on polychoric correlations), as well as the following empirical methods: minimum average partial method^1^, parallel analysis with Glorfeld correction^2^, and the Kaiser rule (# factors = eigenvalues > 1.0). The ESEM was estimated using the mean- and variance-adjusted weighted least-squares (wlsmv) estimator^3, 4^ in Mplus v8.4^5^. Model fit was assessed using the acceptability cutoffs suggested by Hu and Bentler^6^ via the root mean-square error of approximation (RMSEA; acceptable = 0.06), comparative fit index (CFI; acceptable = 0.95), Tucker-Lewis index (TLI; acceptable = 0.95), and standardized root mean-square residual (SRMR; acceptable = 0.08). This model was used to calculate factor scores using the maximum a posteriori (MAP) method. Additional details are provided in the Supplement.

Results from the empirical tests for the number of factors were mixed. Parallel analysis suggested eleven factors or eight principal components; the MAP method suggested five factors; the Kaiser rule suggested eight factors; and the elbow of the scree plot suggested four factors. Based on interpretability and the tendency of empirical methods to over-extract, we settled on five factors, expecting approximately nine items per factor. Fit of the model was acceptable (CFI = 0.96; TLI = 0.95; RMSEA = 0.032 ± 0.001; SRMR = 0.036), and Supplemental Table 1 shows the resulting factor pattern (loading) matrix. Factor 1 (ADHD) comprises all ADHD items and the SIPS Disorganized item (SIPD013) “Trouble with Focus and Attention”; Factor 2 (Psychosis Positive Symptoms) comprises all PRIME screen items; Factor 3 (Mania) comprises all mania items; Factor 4 (Depression) comprises all depression items; and Factor 5 (Psychosis Negative Symptoms) comprises all SIPS Negative items and two Positive items (SIPP090a and SIPP092) “Do people ever tell you that they can't understand you?” and “Disorganized Communication Severity Scale”. Inter-factor correlations (phi) were moderate (mean = 0.40), suggesting a general “p” factor underlying all items^7^.

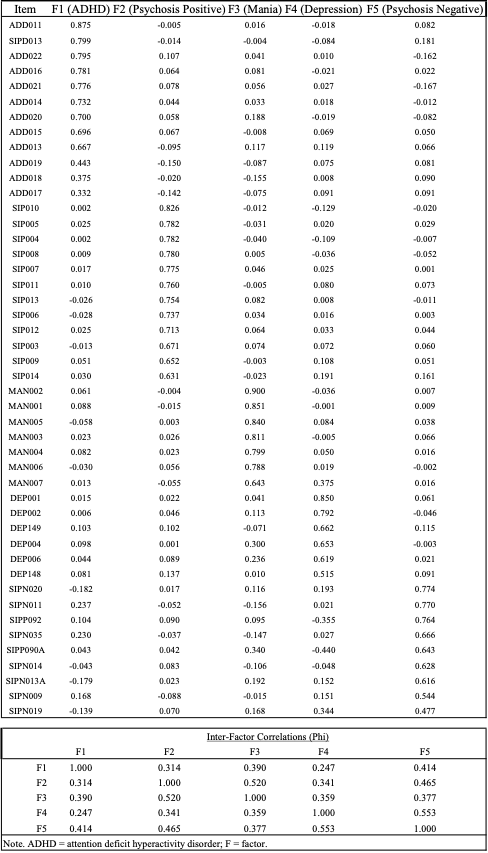

Table 2 Exploratory Structural Equation Model of Clinical Items Overlapping between the GOASSESS and CAPA.

**Supplemental Results**

**Self-reported cannabis use stratified by diagnostic clinical group.**

**Supplemental Figure 1. Self-reported cannabis use rates stratified by clinical diagnosis.**

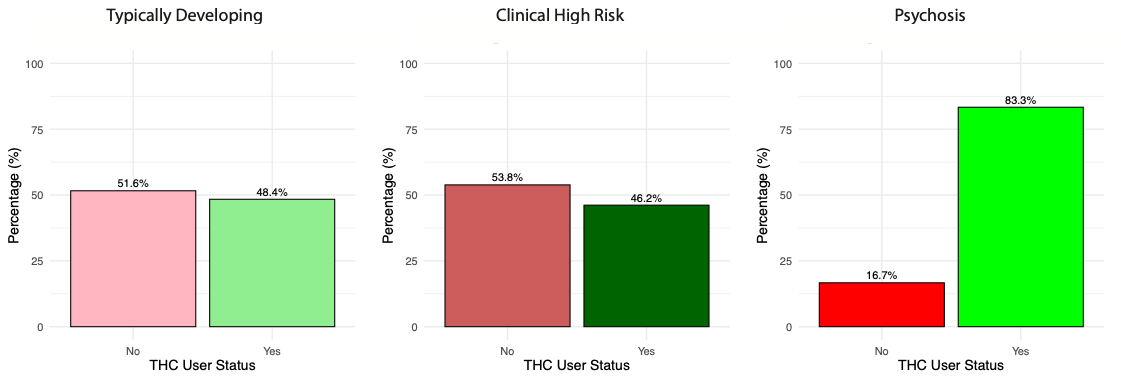

**Supplemental Figure 2. Self-reported frequency of usage stratified by clinical diagnosis**

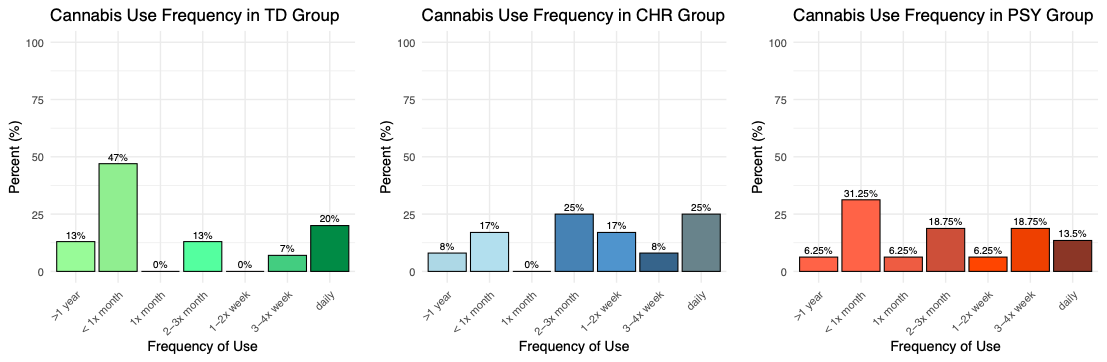

**Dimensional clinical factor scores by diagnosis group**

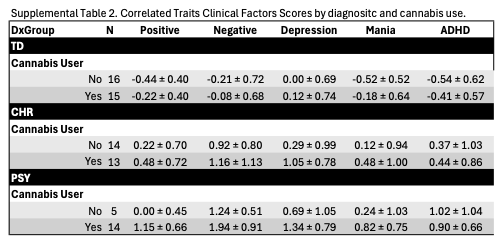

**^1^HMRS glutamate levels are lower in PSY who use cannabis**

As a follow-up to the dimensional analyses, we conducted a sensitivity analysis comparing glutamate levels by diagnostic group while adjusting for age and sex. A linear model revealed a significant main effect of diagnosis on ^1^HMRS glutamate levels, *F*(2, 72) = 4.25, *p* = 0.01 (Supplemental Figure 3). Neither age (*p* = 0.10) nor sex (*p* = 0.52) were significant predictors of glutamate level. Post hoc pairwise comparisons showed that participants with psychosis (PSY) had significantly lower glutamate levels compared to TD (p = 0.01), while comparisons between PSY and CHR (p = 0.15) and between CHR and TD (p = 0.47) were not significant (Supplemental Figure 3).

As a follow up to determine whether the effect of diagnosis group on glutamate levels varied by cannabis use, we stratified the analysis by cannabis user status (Yes/No). Among individuals who reported cannabis use, diagnosis group remained a significant predictor of ^1^HMRS glutamate level, *F*(2, 37) = 3.77, *p* = 0.03, while age (*p* = 0.26) and sex (*p* = 0.74) were not significant (Supplemental Figure 4). In contrast, among non-users, there was no significant effect of diagnosis group on ^1^HMRS glutamate level, *F*(2, 22) = 0.02, *p* = 0.98, while age (*p* = 0.52) and sex (*p* = 0.28) were not significant. Exploratory correlations (Supplemental Results) between glutamate levels and clinical symptoms indicated that higher glutamate levels were associated with lower negative SIPS scores, r(73) = -0.33, *p* = 0.003, which was primarily driven by the PSY cannabis users, r(12) = -0.54. Considered together, these findings indicate robust diagnostic group differences in glutamate levels, independent of demographic covariate and suggest glutamate differences are primarily evident in PSY patients who use cannabis.

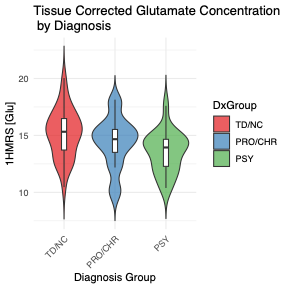

**Supplemental Figure 3. ACC glutamate levels are lower in PSY patients as compared to TD.**

Finally, and as expected, when stratified by clinical diagnosis, symptoms were significantly higher in PSY as compared to CHR and TD in each SIPS domain (positive, negative, disorganized and general) and GAF scores were lower in PSY as compared to CHR and TD (Supplemental Table 1). Given the small sample size of cannabis non-users in the PSY group exploratory pairwise comparisons between cannabis users and non-users were conducted within each diagnostic group to assess differences in clinical symptom scores. Among individuals with PSY, cannabis users exhibited trend level higher positive symptoms, *p* = 0.11, disorganized symptoms (*p* = 0.09) and lower GAF scores (*p* = 0.12), compared to PSY non-users. There were no other trend or significant differences in cannabis users as compared to non-users. CEQ summary scores for paranoia and euphoria did not differ by diagnostic group. There were no diagnostic group differences in age at first cannabis use or length of cannabis use.

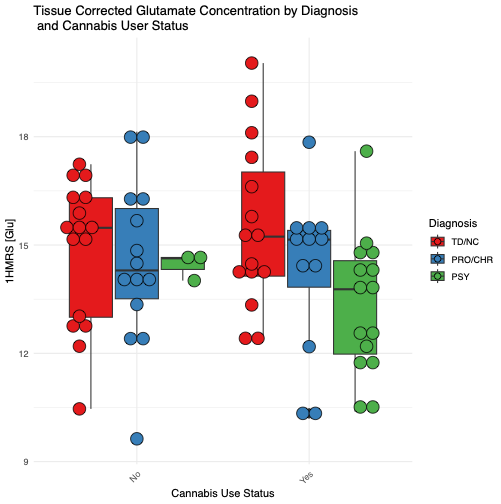

**Supplemental Figure 4. ACC glutamate is lower in PSY patients who use cannabis as compared to TD who use cannabis. Glutamate levels were similar in non-users.**

Cannabis use was more prevalent among participants with psychosis (PSY) than in either the clinical high risk (CHR) or typically developing (TD) groups, consistent with extensive epidemiological evidence linking cannabis exposure to both increased risk for and worsened outcomes in psychosis ^8-10^. In the present sample, 76% of PSY participants reported lifetime cannabis use compared to 48% of CHR and 45% of TD individuals, with planned comparisons revealing a significant PSY–TD difference and a trend toward higher rates in PSY than CHR. These findings are consistent with prior studies that show cannabis use is common in psychosis—up to 40% of patients^10^—and may contribute to illness onset and progression through both direct neurobiological effects of Δ9-THC and shared genetic or environmental vulnerability. The elevated use observed here reinforces the importance of routine cannabis screening in psychosis and early intervention services, especially in the context of rising THC potency and cannabis commercialization^9^.
